# A shift in aggregation avoidance strategy marks a long-term direction to protein evolution

**DOI:** 10.1101/176867

**Authors:** S.G. Foy, B.A. Wilson, J. Bertram, M.H.J. Cordes, J. Masel

## Abstract

To detect a direction to evolution, without the pitfalls of reconstructing ancestral states, we need to compare “more evolved” to “less evolved” entities. But because all extant species have the same common ancestor, none are chronologically more evolved than any other. However, different gene families were born at different times, allowing us to compare young protein-coding genes to those that are older and hence have been evolving for longer. To be retained during evolution, a protein must not only have a function, but must also avoid toxic dysfunction such as protein aggregation. There is conflict between the two requirements; hydrophobic amino acids form the cores of protein folds, but also promote aggregation. Young genes avoid strongly hydrophobic amino acids, which is presumably the simplest solution to the aggregation problem. Here we show that young genes’ few hydrophobic residues are clustered near one another along the primary sequence, presumably to assist folding. The higher aggregation risk created by the higher hydrophobicity of older genes is counteracted by more subtle effects in the ordering of the amino acids, including a reduction in the clustering of hydrophobic residues until they eventually become more interspersed than if distributed randomly. This interspersion has previously been reported to be a general property of proteins, but here we find that it is restricted to old genes. Quantitatively, the index of dispersion delineates a gradual trend, i.e. a decrease in the clustering of hydrophobic amino acids over billions of years.

## Introduction

Proteins need to do two things to ensure their evolutionary persistence: fold into a functional conformation whose structure and/or activity benefit the organism, and also avoid folding into harmful conformations. Amyloid aggregates are a generic structural form of any polypeptide, and so pose a danger for all proteins (Monsellier and Chiti 2007). Several lines of evidence suggest that aggregation avoidance is a critical constraint during protein evolution. Highly expressed genes are less aggregation-prone (Tartaglia *et al.* 2007), and evolve more slowly due to greater selective constraint against alleles that increase the proportion of mistranslated variants that misfold (Drummond *et al.* 2005; Drummond and Wilke 2008). Genes that homo-oligomerize or are essential (Chen and Dokholyan 2008) or that degrade slowly (De Baets *et al.* 2011) are also less aggregation-prone. Aggregation-prone stretches of amino acids tend to have translationally optimal codons (Lee *et al.* 2010), and be flanked by “gatekeeper” residues (Rousseau *et al.* 2006). Disease mutations are enriched for aggregation-promoting changes (Reumers *et al.* 2009; De Baets *et al.* 2015), and known aggregation-promoting patterns are underrepresented in natural protein sequences (Broome and Hecht 2000; Buck *et al.* 2013). Thermophiles, whose amino acids need to be more hydrophobic, show exaggerated aggregation-avoidance patterns (Thangakani *et al.* 2012).

Here we ask whether and how proteins get better at avoiding aggregation during the course of evolution. Absent a fossil record or a time machine, biases introduced during the inference of ancestral protein states (Williams *et al.* 2006; Trudeau *et al.* 2016) make it difficult to assess how past proteins systematically differed from their modern descendants. We have therefore developed an alternative method to study protein properties as a function of evolutionary age, one that does not rely on ancestral sequence reconstruction.

While all living species share a common ancestor, all proteins do not. It has become clear that protein-coding genes are not all derived by gene duplication and divergence from ancient ancestors, but instead continue to originate de novo from non-coding sequences (McLysaght and Guerzoni 2015). Different gene families (i.e. sets of homologous genes) therefore have different ages, and the properties of a gene can be a function of age.

The age of a gene can be estimated by means of its “phylostratum”, which is defined by the basal phylogenetic node shared with the most distantly related species in which a homolog of the gene in question can be found (Domazet-Lošo *et al.* 2007). Failure to find a still more distantly related protein homolog (i.e. failure of a gene to appear older) can have multiple causes. First, more distantly related homologs might not exist, as a consequence of de novo gene birth either from intergenic sequences or from the alternative reading frame of a different protein-coding gene (the latter yielding nucleotide but not amino acid homology). Second, apparent age might indicate the time not of de novo birth but of horizontal gene transfer (HGT) from a taxon for which no homologous genes have yet been sequenced. Third, independent loss of the entire gene family in multiple distantly related lineages can yield a pattern of apparent gain. Fourth, divergence between gene duplicates might be so extreme that homology can no longer be detected.

The diversity of sequenced taxa now available makes the second possibility (HGT) increasingly unlikely, especially outside microbial taxa that experience high levels of HGT; here we minimize this possibility by focusing on the set of mouse genes. The same wealth of sequenced taxa also makes the third possibility (phylogenetically independent loss of the entire gene family) unlikely, given the large number of independent loss events implied. More importantly, neither HGT nor independent loss are likely to drive systematic trends in protein properties as a function of apparent gene age; instead, they are likely to dilute any underlying patterns resulting from other determinants of apparent gene age.

Most critiques of the interpretation of phylostratigraphy in de novo gene terms therefore focus on the fourth possibility, specifically the concern that trends may be driven by biases in the degree to which homology is detectable (Albà and Castresana 2007; Moyers and Zhang 2015; Moyers and Zhang 2016; Moyers and Zhang 2017). In particular, homology is harder to detect for shorter and faster-evolving proteins, which might therefore appear to be young, giving false support to the conclusion than young genes are shorter and faster-evolving. The problem of homology detection bias extends to any trait that is correlated with primary factors, such as length or evolutionary rate, that directly affect homology detection. We previously studied such a trait, intrinsic structural disorder (ISD), and found that statistically correcting for evolutionary rate did not affect the results, and that statistically correcting for length made them stronger (Wilson *et al.* 2017). This suggested that the pattern in ISD was likely driven by time since de novo gene birth, rather than by homology detection bias.

Here we trace a number of other protein properties as a function of apparent gene family age, including aggregation propensity and hydrophobicity, and find a particularly striking trend for the degree to which hydrophobic residues are clustered along the primary sequence. This trend, as with the previous ISD work, experiences negligible change after correction for length, evolutionary rate, and expression, and is thus not a result of homology detection bias. Our results point to a systematic shift in the strategies used by proteins to avoid aggregation, as a function of the amount of evolutionary time for which they have been evolving.

## Results

We assigned mouse genes to gene families and to times of origin, and assigned a protein aggregation propensity score to each protein on the basis of its amino acid sequence (see Methods). No clear trend is seen in aggregation propensity as a function of gene age (Fig. 1), although all genes (black) show lower aggregation propensity than would be expected if intergenic mouse sequences were translated into polypeptides (blue). Note that intergenic sequences represent not only the raw material from which de novo genes could emerge, but also the fate of any sequence, e.g. a horizontally transferred gene, that is subjected to neutral mutational processes.

**Fig. 1.**
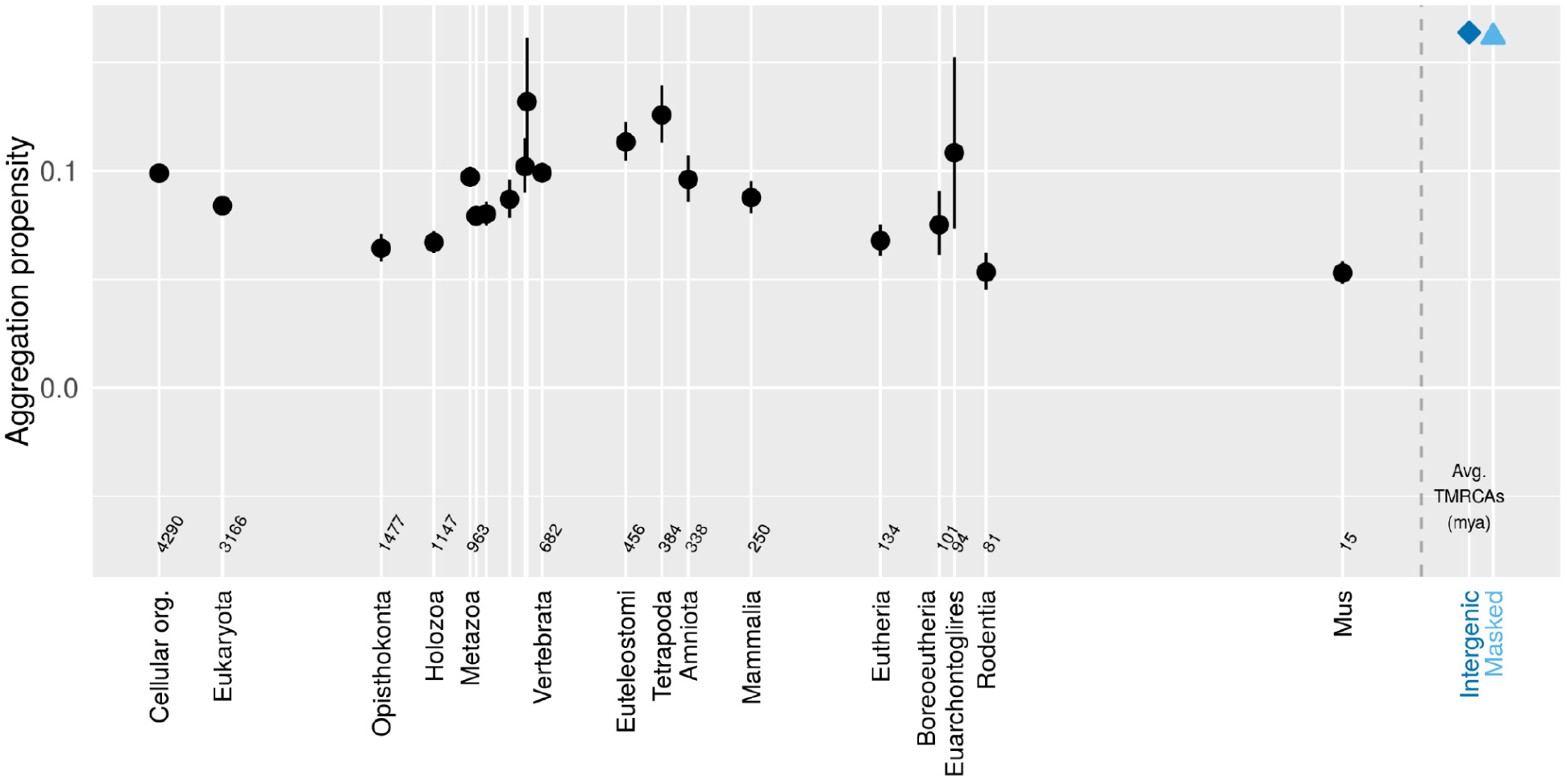
Mouse genes show little pattern in aggregation propensity (assessed via TANGO) as a function of age. Genes (black) show less aggregation propensity than intergenic controls (blue). Back-transformed central tendency estimates +/- one standard error come from a linear mixed model applied to transformed data, where gene family and phylostratum are random and fixed terms respectively. Importantly, this means that we do not treat genes as independent data points, but instead take into account phylogenetic confounding, and use gene families as independent data points. Times to most recent common ancestor (TMRCAs) for most phylostrata were taken from TimeTree.org (Kumar *et al.* 2017) on February 18, 2016 and that for *M. pahari* was taken May 7, 2018. We used the arithmetic means of the TMRCAs of the focal taxon shown on the x-axis and the preceding taxon (i.e. the estimated midpoint of the interior branch of the tree). Cellular organism age is shown as the midpoint of the last universal common ancestor and the last eukaryotic common ancestor. Taxon names, some of which are omitted for space reasons, follow the sequence Metazoa, Eumetazoa, Bilateria, Deuterostomia, Chordata, Olfactores, Vertebrata, Euteleostomi, Tetrapoda, Amniota, Mammalia, Eutheria, Boreoeutheria, Euarchontoglires, Rodentia, Mus. The grey dashed line shows the 0 time, with control sequences to the right of it.

However, striking patterns emerge when we decompose aggregation avoidance into the effect of amino acid composition (with hydrophobic amino acids making aggregation more likely), and the effect of the exact order of a given set of amino acids. The contribution of amino acid composition alone can be assessed by scrambling the order of the amino acids (Fig. 2, top), revealing that young genes make greater use of amino acid composition to avoid aggregation. The pattern is mirrored by other measurements of the hydrophobicity of the amino acid composition (Fig. 2, middle panels on the fraction of hydrophobic residues and on intrinsic structural disorder, the latter previously reported by Wilson *et al.* (2017)), with an increase in hydrophobicity taking place over ^~^200-400 million years. Previously reported differences in the aggregation propensity (Tartaglia *et al.* 2005) and hydrophobicity (Mannige *et al.* 2012) of proteomes from different organisms might therefore be accounted for by systematic variation among species in the composition of old vs. young genes; in our analysis, all proteins were taken from the same mouse species, removing this confounding factor. Analyses focused on a set of ancestral reconstructed sites also find a trend of recently increasing hydrophobicity in Drosophilid genomes (Yampolsky and Bouzinier 2010) that is ongoing even for ancient gene families (Yampolsky *et al.* 2017), although this data is subject to the bias of observing slightly deleterious substitutions more often than the reverse (Hurst *et al.* 2006; McDonald 2006).

**Fig. 2.**
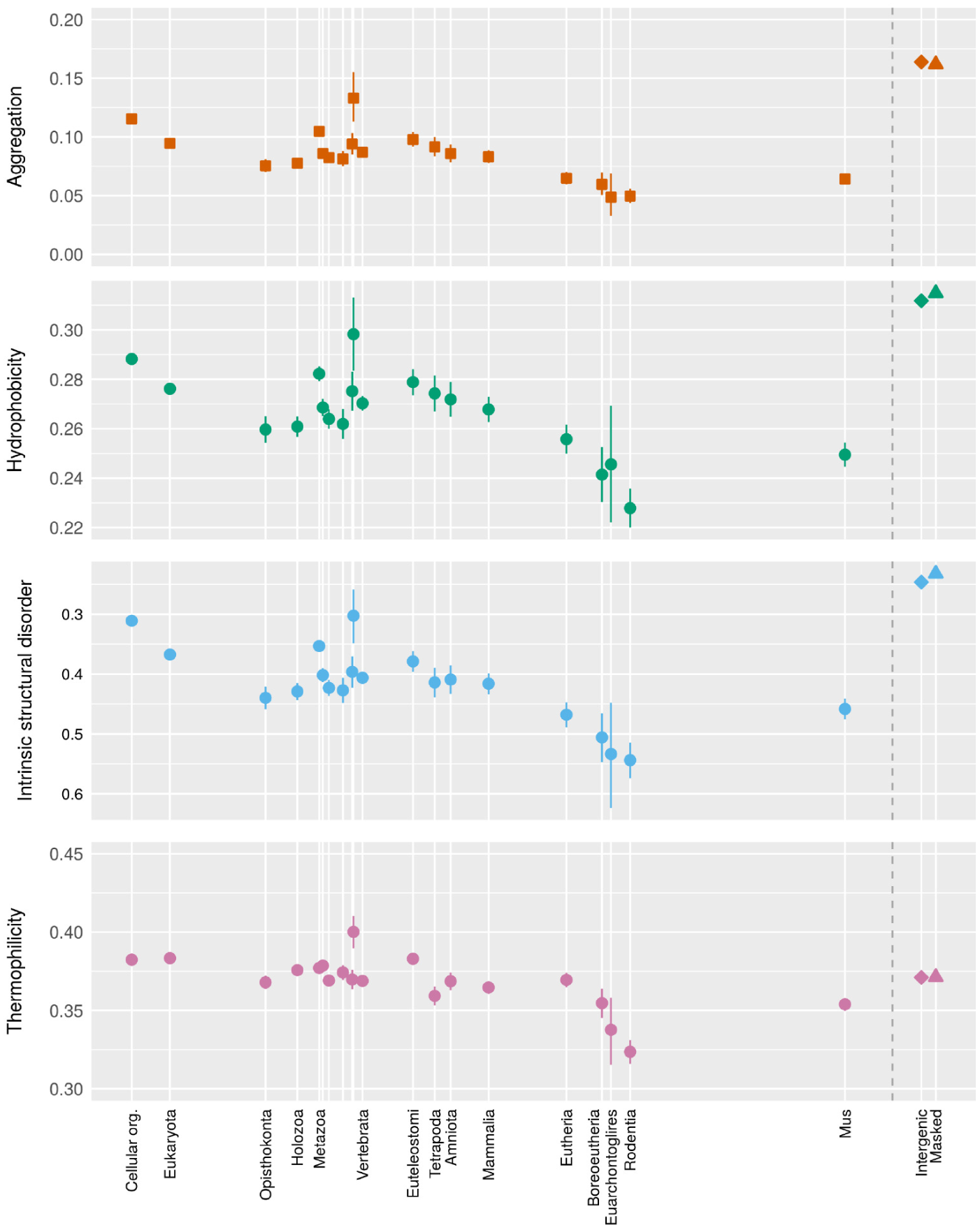
Four different measures for the hydrophobicity of the amino acid content as a function of gene family age. “Aggregation” represents the average TANGO results from 50 scrambled versions of each gene, and hence captures the effect of amino acid composition on TANGO’s estimate of β-aggregation propensity. The use of scrambled genes is indicated by squares, with unscrambled genes as circles and intergenic controls as diamonds or triangles depending on whether repeat sequences are excluded. Hydrophobicity gives the fraction of amino acids that are FLIMVW. The “oiliness” measurement ofMannige et al. (2012), namely content of FLIV, is similar. Intrinsic structural disorder scores are as previously reported inWilson et al. (2017), shown here for more phylostrata, and inverted for easier comparison with other metrics. Thermophilicity represents the content of ILVYWRE, as analyzed byBoussau et al. (2008), subjected to a Box-Cox transform with λ= 2.412 prior to model fitting; thermophilicity is dominated by the same general hydrophobicity trend as the other measures. While the trend as a function of gene age is similar in each case, the aggregation measurement shows the most striking deviation from intergenic control sequences. Back-transformed central tendency estimates +/- one standard error come from a linear mixed model, where gene family and phylostratum are random and fixed terms respectively; λ=0.93 is used for hydrophobicity, other transforms are described in the Methods. The x-axis is the same as for Figure 1.

The contribution of amino acid ordering alone, independent from amino acid composition, can be assessed as the difference between the aggregation propensity of the actual protein and that of a scrambled version of the protein. We expected real proteins to be less aggregation-prone than their scrambled controls (Buck *et al.* 2013), and confirmed this for the very oldest proteins (Fig. 3, orange confidence intervals for genes shared with prokaryotes lie below 0). But surprisingly, the opposite was true for young genes (Fig. 3, orange values for phylostrata from metazoa onward lie above 0). In other words, they are more aggregation-prone than would be expected from their amino acid composition alone.

**Fig. 3.**
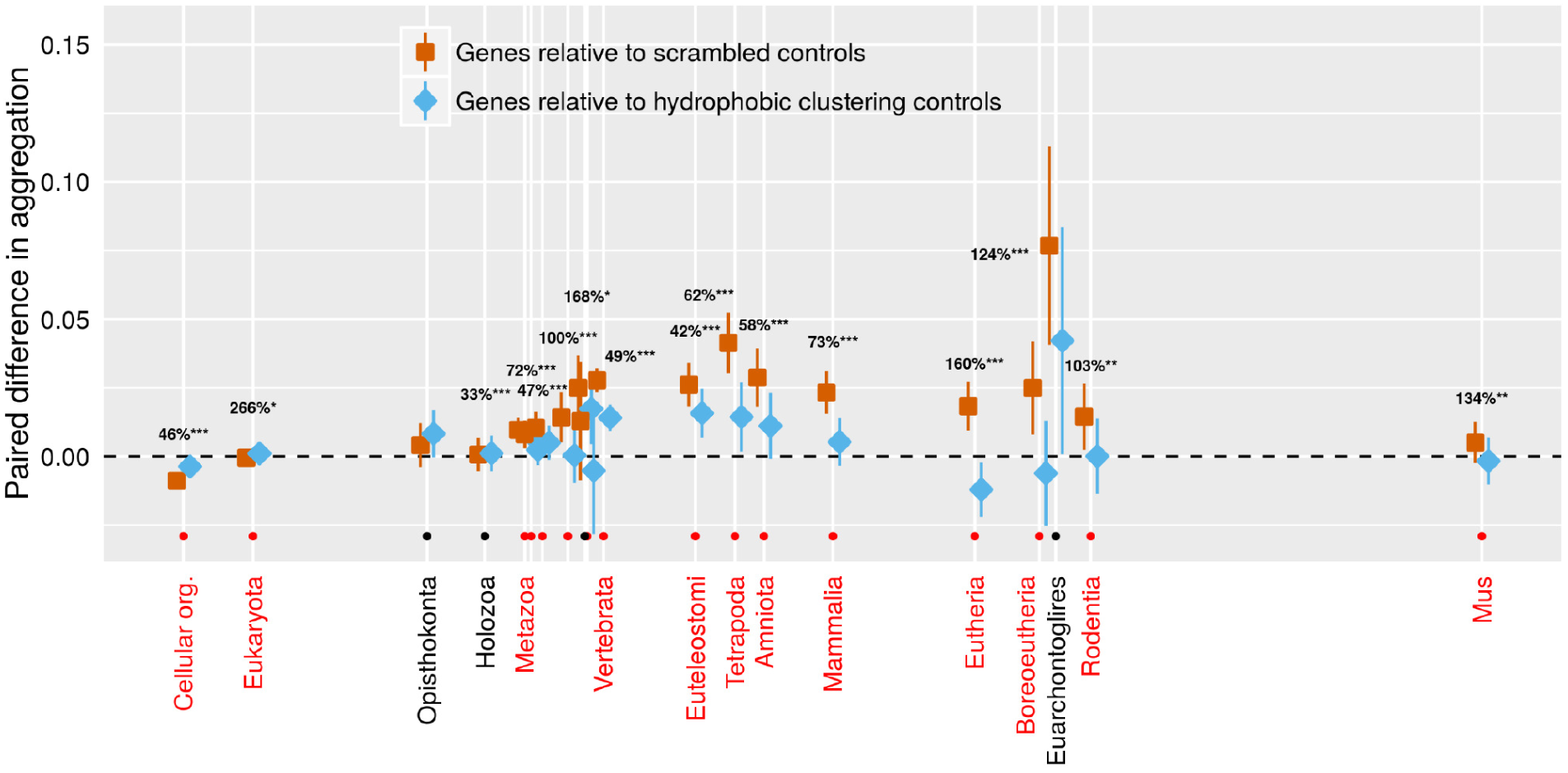
Only very old genes have aggregation propensities lower than that expected from their amino acid composition alone (orange < dashed line expectation of 0). This puzzling finding is reduced when we account for clustering (blue is closer than orange is to the 0 dashed line) using a scrambled sequence that is controlled to have a similar clustering value. The clustering of hydrophobic amino acids in young genes acts to increase their aggregation propensity. 95% confidence intervals are shown, based on a linear mixed model where gene family and phylostratum are random and fixed terms respectively. Note that blue and orange confidence intervals should be compared only to the reference value of zero, and not to each other, due to the paired nature of the data. For phylostrata shown in red and indicated by an orange dot, the difference between blue and orange was significant (\**p*<0.01, \*\**p*<0.001, \*\*\**p*<0.0001), and the percentage of deviation from 0 accounted for by the control is shown. For most phylostrata where the difference between blue and orange was non-significant (indicated by a black dot and black text), the orange deviated little from 0, so there was little or nothing for the blue clustering control to account for. Results are shown for TANGO; results for Waltz trend in the same direction but are weaker (Fig. S5). Orange values come from the mean of 50 scrambled sequences per gene, blue from a single scrambled sequence with a closely matched clustering value. The x-axis is the same as for Figure 1.

One possible source of increased aggregation propensity is if young genes, struggling to achieve any kind of fold at all given their low hydrophobicity (Dill 1990), cluster their few hydrophobic amino acid residues closer together along the sequence. Such clustering could allow proteins to evolve small, foldable, potentially functional domains within an otherwise disordered sequence (Uversky *et al.* 2000). Alternatively and still more primitively, very highly localized clustering could produce short peptide motifs that cannot fold independently but acquire structure conditionally through binding or oligomerization (Gunasekaran *et al.* 2004; Davey *et al.* 2012). Hydrophobic clustering also increases the danger of aggregation (Monsellier *et al.* 2007); indeed, there is significant congruence between mutations that increase the stability of a fold and those that increase the stability of the aggregated or otherwise misfolded form (Sánchez *et al.* 2006).

We find that young genes do show hydrophobic clustering, while very old genes show interspersion of hydrophobic amino acid residues (Fig. 4), and that this accounts for much of the excess aggregation propensity of young genes relative to scrambled controls (Fig. 3 blue points are closer to zero than orange points). Previous reports have suggested that the danger of aggregation selects against hydrophobic clustering (Monsellier *et al.* 2007). In other words, among consecutive blocks of amino acids, the variance in hydrophobicity is lower than the mean, i.e. the index of dispersion is less than one in proteins overall (Irbäck *et al.* 1996; Schwartz *et al.* 2001) and in the core of protein folds (Patki *et al.* 2006). In the present analysis, this holds true only for old, highly evolved proteins. Younger proteins not only appear less evolutionarily constrained to intersperse polar and hydrophobic residues, but to the contrary, their hydrophobic residues show excess concentration near one another along the sequence, increasing aggregation propensity. Our results are extremely robust when we control for protein length, evolutionary rate, and expression level (Fig. S1). Similar results, albeit not extending quite as far back in time, are found using the normalized mean length of runs of hydrophobic amino acid lengths FLIMVW (Fig. S2) as by using the more sophisticated published metric of the degree to which these amino acids are clustered (Irbäck *et al.* 1996; Irbäck and Sandelin 2000) shown in Fig. 4.

**Fig. 4.**
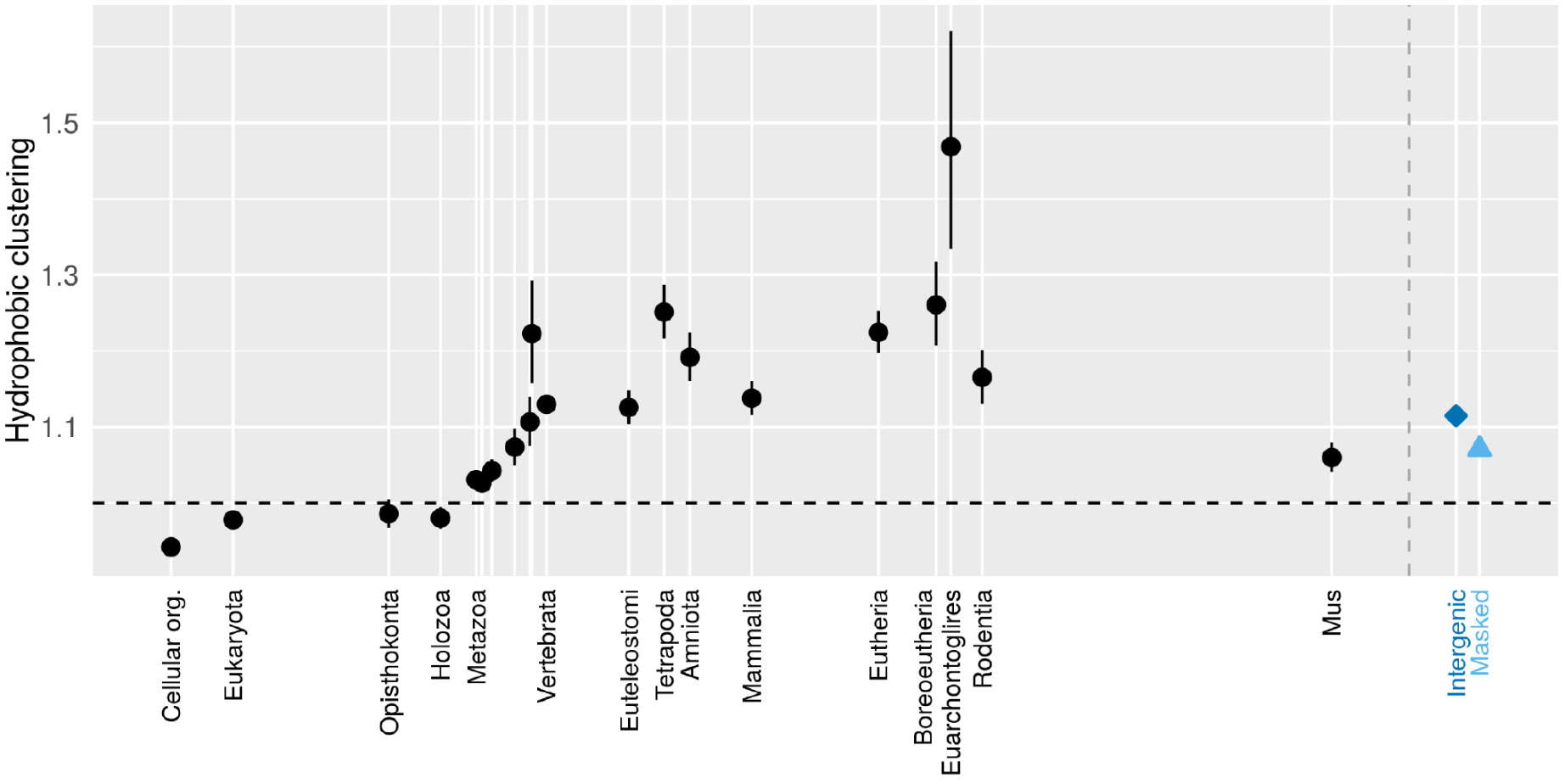
Clustering initially follows that of its raw material, and evolves rapidly upward at first, but then decays downward extremely slowly, indicating a long-term direction of evolution. Only the oldest genes have hydrophobic amino acids spread out from each other, as previously reported; young genes have clustered hydrophobic amino acids. Back-transformed central tendency estimates +/- one standard error come from a linear mixed model, where gene family and phylostratum are random and fixed terms respectively. The x-axis is the same as for Figure 1.

We investigated whether the difference might be explained by differences in the frequencies of transmembrane proteins as a function of gene age. Unfortunately, sequence-based prediction of transmembrane status is likely directly confounded with clustering, and only 137 mouse proteins have been experimentally verified as transmembrane. The increased clustering of transmembrane proteins was barely significant as a fixed effect within our linear model (*p*=0.042). Because annotations are available for so few genes, we do not know whether transmembrane proteins are more likely to be old or young, and hence whether transmembrane status helps explain the trend or make it still more puzzling. The weak effect size suggests that transmembrane status might explain at best only a small portion of the trend.

We checked whether this trend in clustering is also found in the proteins of *Saccharomyces cerevisiae* (Fig. S3), which is the other species for which homologous gene family annotation was combined with gene age annotation (Wilson *et al.* 2017). The very youngest 499 putative gene families (unique to *S. cerevisiae*, and which might therefore contain non-coding sequences annotated in error, although to minimize this problem, genes annotated as “dubious” are excluded) had a clustering value of 1.035 (66% CI 1.024-1.047; central tendency and CI backtransformed from the central tendency estimate +/- one standard error derived from a linear model with and gene family as a random effect). The oldest 1,966 gene families (with homologs in prokaryotes) had clustering 0.890 (66% CI 0.886-0.895), even lower than clustering of 0.943 (66% CI 0.939-0.946) found in mouse gene families of the same age. Among the 2,467 gene families allocated to eight phylostrata of intermediate age, we found no significant differences among the phylostrata (p=0.6, likelihood ratio test of linear model with gene family and random effect and phylostratum as putative fixed effect), which range from genes shared only with *S. paradoxus* to genes shared with distantly related eukaryotes. The clustering in all these phylostrata was lower than we expected from our mouse results, at 0.951 (66% CI 0.945- 0.958). These results, shown in Figure S3, suggest that low clustering evolves far more rapidly, at least in the earlier stages, in unicellular yeast with short generation times and large population sizes than it does in the ancestral lineage of mice. However just as for the mouse lineage, saturation is not reached for gene families dating back “only” to an early eukaryote; genes with prokaryotic homologs have even lower clustering values than those with homologs in distantly related eukaryotes but not prokaryotes.

Clustering is a metric for which genes that have been evolving for longer have different properties from genes that are “less evolved”. There must either be a long-term trend in the clustering values of newborn genes as a function of the time at which they are born, or else there has been a long-term direction to evolution over billions of years. We consider the latter possibility more plausible than the former. This directionality of evolution can be interpreted as a slow shift from a primitive strategy for avoiding misfolding in young genes to more subtle strategies in old genes.

The primitive aggregation avoidance strategy used by young genes is simply to avoid the most hydrophobic amino acids (Fig. 2), creating intrinsic structural disorder (Linding *et al.* 2004; Thangakani *et al.* 2012; Banerjee and Chakraborty 2017; Wilson *et al.* 2017). Given such an amino acid composition, young genes might form an early folding nucleus by concentrating hydrophobic amino acids in localized regions of the sequence (Fig. 4, right), while still keeping total hydrophobicity and hence aggregation propensity within tolerable limits (Figs. 1-2). Such a folding nucleus would not necessarily be an entire independently folded domain. In particular, some origin theories posit that ancient proteins first achieved folding by becoming structured only upon binding to some interaction partner (Soding and Lupas 2003; Zhu *et al.* 2016). In contemporary proteins, potential representatives of nascent structure are found in intrinsically disordered proteins that contain peptide-length binding motifs (small linear interaction motifs; SLiMs), many of which become ordered when bound to a partner (Davey *et al.* 2012). We do not, however, find that young genes have more known SLiMs (Fig. S4).

In contrast to young genes, older genes have higher hydrophobicity, which must be offset by the evolution of other aggregation-avoidance strategies (Thangakani *et al.* 2012). For such changes to occur through descent with modification probably happens only slowly. Under the assumption that amino acid composition at birth does not vary systematically as a function of the time of birth, we could conclude that changing the amino acid composition of a protein takes ~200-400 million years (Fig. 2). In contrast, changing the index of dispersion might require such a large number of changes that it is extraordinarily slower, with a consistent direction to evolution visible over the entire history of life back to our common ancestor with prokaryotes.

Note that our two youngest phylostrata, the Mus phylostratum of *Mus musculus* genes shared only with *M. pahari*, and the Rattus phylostratum of *M. musculus* genes shared with rats, show less clustering than other young genes, suggesting that rapid change in the index of dispersion may be possible (in the other direction) after all, on short and recent timescales. However, very young gene families are subject to significantly higher death rates than other gene families (Palmieri *et al.* 2014). With gene family loss so common at first, it is possible that the rapid initial increase in clustering is due to differential retention of gene families with highly clustered amino acids. This interpretation of the data is consistent with explaining how slow the later fall in clustering is, by positing that descent with modification is constrained to change clustering values slowly.

The youngest genes show similar clustering to what would be expected were intergenic sequences to be translated (Fig. 4, blue). Clustering of amino acids translated from non-coding intergenic sequences is a direct consequence of the clustering of nucleotides; indices of dispersion at the nucleotide level are all above the expectation of one from a Poisson process, in the range 1.2-1.9 for intergenic sequences and 1.1-1.8 for masked intergenic sequences, depending on which nucleotides are considered. (The lowest indices are found for the GC vs. AT contrast, presumably due to avoidance of CpG sites causing a general paucity of clusters of G and C.) Very short tandem duplications, e.g. as may arise from DNA polymerase slippage, automatically create segments in which the duplicated nucleotide is overrepresented; observed nucleotide clustering values greater than one can therefore be interpreted as a natural consequence of mutational processes. The consequence of this mutational pattern is therefore a small and fortuitous degree of preadaptation, i.e. intergenic sequences have a systematic tendency toward higher clustering than “random”, in a manner that facilitates the de novo birth of new genes.

## Discussion

As discussed in the Introduction, apparent gene family age can be a function of time since i) gene birth, ii) HGT, iii) divergence from other phylogenetic branches all of which have independently lost all members of the gene family, or iv) rapid divergence of a gene made homology undetectable. In all cases, our results describe evolutionary outcomes as a function of time elapsed since that event. In the case of our primary result on clustering, this means that genes appear with clustering values similar to those expected from intergenic sequences, are retained only if their clustering is exceptionally high, and then show gradual declines in clustering after that.

We believe that gene birth is the most plausible driver of our results. HGT is rare in more recent ancestors of mice, simultaneous loss in so many branches is unlikely, and statistical correction for evolutionary rate, length and expression (Fig. S1) has, in contradiction to the predictions of homology detection bias, a negligible effect on our results. However, our results on the evolution of protein properties following a defining event remain of interest under all scenarios of what the gene-age-determining event is.

There are three ways to explain subsequent patterns as a function of gene family age. The two mentioned so far are biases in retention after birth, and descent with modification. The third possibility is that the conditions of life were significantly different at different times, and hence so were the biochemical properties of proteins born/transferred/rapidly diverged at that time. Specifically, ancestral sequence reconstruction techniques have been used to infer that proteins in our ancestral lineage became progressively less thermophilic (Gaucher *et al.* 2008). This might explain why young genes have fewer strongly hydrophobic amino acids; they were born at more permissive lower temperatures. However, ancestral reconstruction techniques are likely biased toward consensus amino acids that are fold-stabilizing (Steipe *et al.* 1994; Lehmann *et al.* 2000; Godoy-Ruiz *et al.* 2004; Bloom and Glassman 2009) and hence may be more hydrophobic (Williams *et al.* 2006; Trudeau *et al.* 2016). Alarmingly, ancestral reconstruction also suggests that the ancestral mammal was a thermophile (Trudeau *et al.* 2016), although Drosopholid reconstructions are compatible with a trend in the opposite direction to reconstruction bias, towards greater hydrophobicity with time (Yampolsky and Bouzinier 2010; Yampolsky *et al.* 2017).

The main trend that we see of hydrophobicity/thermophilicity as a function of gene age is on shorter timescales; for older gene families, billions of years of common evolution has erased the differences in starting points. It is the more subtle signal of hydrophobic amino acid interspersion that shows the long-term pattern in our analysis. However, variation in the conditions of life at the time of gene origin remains a plausible explanation for the idiosyncratic differences between phylostrata, i.e. for the remaining, statistically meaningful deviations of individual phylostrata from the trends reported here.

We have already invoked differential retention as a possible driver of the short-term evolutionary increase in the clustering values of young genes. It is logically possible that the long-term trend in clustering values is also a result of differential retention; if gene families with higher clustering values are more likely to be lost, different gene ages represent different spans of time in which this loss has had an opportunity to occur. Given the billion year time scales and thus enormous number of lost gene families this implies, this seems at present a less plausible scenario than descent with modification for different durations following different dates of origin. In other words, descent with modification seems the most plausible of the three possible drivers of biochemical patterns as a function of gene age, independently of what exactly “gene age” means.

Note that our findings go in the opposite direction to those of Mannige et al. (2012), who used more speciation-dense branches as a proxy for longer effective evolutionary time intervals, to infer an evolutionary trend away from, rather than toward, hydrophobicity. Part of this discrepancy may arise from differences in which proteins are present in which species, which could be a confounding factor when Mannige et al. attributed proteome-wide trends to descent with modification. Mannige et al. also confirmed their results for single genes, but did not, in that portion of their analysis, also confirm that results were not sensitive to the difficulty of scoring speciation-density in prokaryotes.

We propose that our findings may be best explained by three phases of protein evolution under selection for proteins that both avoid misfolding and have a function. First, a filter during the gene birth process gives rise to low hydrophobicity in newborn genes (Wilson *et al.* 2017), as the simplest way to avoid misfolding. Second, young genes with their few hydrophobic amino acids clustered together are more likely to have functional folds that remain adaptive for some time after birth, and so are differentially retained in the period immediately after birth (when young genes are subject to very high rates of attrition (Palmieri *et al.* 2014)). Finally these two initial trends are both slowly reversed by descent with modification, continuing over billions of years of evolutionary search for better solutions for exceptions to the intrinsic correlation between propensity to fold and propensity to misfold.

The protein folding problem is notoriously hard. Here we see that it isn’t just hard for human biochemists – it’s so hard that evolution struggles with it too. Proteins evolve to find stable folds despite the correlated and ever-present danger of aggregation. They do so via a slow exploration of an enormous sequence space, a search that has yet to saturate after billions of years (Povolotskaya and Kondrashov 2010). Given the enormous space that has already been searched, existing protein folds, especially of older gene families, may therefore be a highly unrepresentative sample of the typical behaviors of polypeptide chains. Protein folds are best thought of as a collection of corner cases and idiosyncratic exceptions, which are hard to find even for evolution, let alone for our “free-modeling” techniques to predict ab initio.

## Methods

*M. musculus* proteins from Ensembl (v73) were assigned gene families and gene ages as described elsewhere (Wilson *et al.* 2017). To briefly outline this previous procedure, BLASTp (Altschul *et al.* 1997) against the National Center for Biotechnology Information (NCBI) nr database with an E-value threshold of 0.001 was used for preliminary age assignments for each gene, followed by a variety of quality filters. Genes unique to one species were excluded because of the danger that they were falsely annotated as protein-coding genes (McLysaght and Hurst 2016), leaving Rodentia as the youngest phylostratum. Paralogous genes were clustered into gene families, and a single age was reconciled per gene family, which filtered out some inconsistent performance of BLASTp. Numbers of genes and gene families in each phylostratum can be found for mouse in Table S1 of Wilson et al. (2017). “Cellular Organisms” contains all mouse gene families that share homology with a prokaryote. Yeast gene family and phylostratum annotation is taken from Table S7 of Wilson et al. (2017).

For greater resolution at shorter timescales, we used the recently sequenced *M. pahari* genome (Thybert *et al.* 2018) to compile a younger phylostratum, using Ensembl’s orthology annotation (Herrero *et al.* 2016) to find homologs in *M. musculus*. Of the 789 putative proteins excluded in Wilson et al. (2017) as being unique to *M. musculus*, 155 also had homologs in *M. pahari*. 9 of these also had Ensembl ortholog assignments among members of older gene families, and were excluded. BLASTp detected only one pair hitting each other among the genes with e-value < 0.001; these were placed together while each of the others was placed in its own gene family, collectively forming the youngest phylostratum to be analyzed. Note also that Ensembl ortholog annotation is not as rigorous a filter to remove false positives as the rat vs mouse dN/dS measures used by Wilson et al. (2017) for older phylostrata. We therefore do not expect this youngest Mus phylostratum to be entirely free of false positives. This likely explains why its hydrophobicity metrics are lower than those of Rattus. The fact that hydrophobicity is still significantly elevated above that of controls (especially as measured by ISD and by predicted aggregation propensity of scrambled sequences) suggests that the problem of contamination with sequences that are not protein-coding genes is not so profound as to exclude the phylostratum. However, it should be interpreted with caution.

Intergenic control sequences were also taken from previous work (Wilson *et al.* 2017). Briefly, one intergenic control sequence per gene was taken 100nt downstream from the end of the 3’ end of the transcript, with stop codons excised until a length match to the neighboring protein-coding gene was obtained. A second control sequence per gene began 100nt further downstream. This choice of location ensures that control sequences are representative of genomic regions in which protein-coding genes are found. One version of the control sequences used all intergenic sequences for this procedure, a second used only RepeatMasked (Smit *et al.* 2015) intergenic sequences.

Aggregation propensity was scored using TANGO (Fernandez-Escamilla *et al.* 2004) and Waltz (Maurer-Stroh *et al.* 2010). We counted the number of amino acids contained within runs of at least five consecutive amino acids scored to have >5% aggregation propensity, added 0.5, and divided by protein length to obtain a measure of the density of aggregation-prone regions. For those scores derived using TANGO, we then performed a Box-Cox transformation (λ=0.362, optimized using only coding genes not controls, Q-Q plot shown in Fig. S6A) prior to linear model analysis in Figs. 1 and S1. Box-Cox λ values were determined using maximum-likelihood estimation (Box and Cox 1964) as implemented in geoR (https://CRAN.R-project.org/package=geoR). Central tendency estimates and confidence intervals derived from these models were then back transformed for the plots. Paired differences in TANGO scores or Waltz scores between genes and scrambled controls were not transformed. Results were qualitatively indistinguishable when runs of at least six consecutive amino acids were analyzed instead of runs of at least five.

“Clustering” was assessed as a normalized index of dispersion, i.e. by comparing the variance in hydrophobicity between blocks of consecutive amino acids to the mean hydrophobicity (Irbäck *et al.* 1996). Examples of high and low clustering are shown in Fig. 5. We used *s* = 6, with different values of *s* yielding qualitatively similar results. Where the amino acid length was not divisible by six, a few amino acids were neglected at one or both ends, yielding a truncated length of *N*, and we used the average clustering measure *ψ* across different phases for the blocking procedure. In Fig. 4, we average over all phases using the maximum number of blocks, e.g. only one phase for values of *N* divisible by 6. Results when we average over all 6 phases are very similar. Following past practice, we transformed amino acid sequences into binary hydrophobicity strings by taking the six amino acids FLIMVW as hydrophobic (+1) and scoring all the other amino acids as −1. We summed hydrophobicity scores to a value *σ_k_* for each block *k* = 1, …, *N*/*s* and 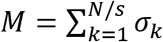 overall (Irbäck and Sandelin 2000). Our clustering score is a normalized index of dispersion

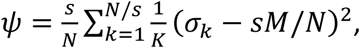

where the normalization factor for length *N* and total hydrophobicity *M* of a protein is

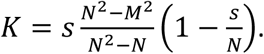

**Fig. 5.**
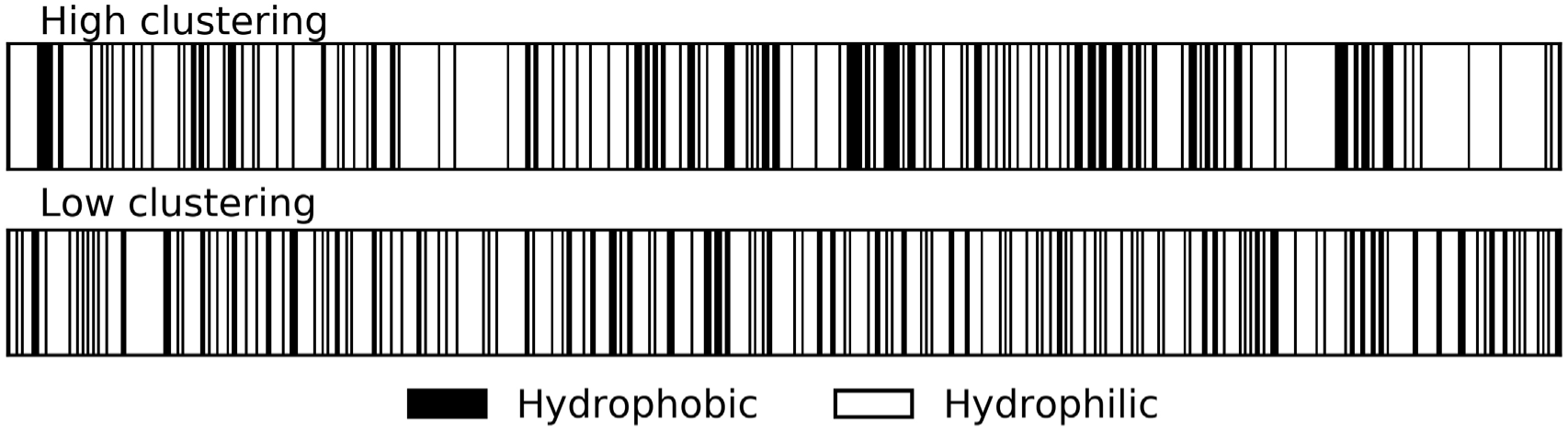
Illustration of the distribution of hydrophobic residues along the primary sequence of proteins with high vs. low clustering, of similar lengths and net hydrophobicities. The high clustering gene Fzd5 has length 585 amino acids, 31.5% hydrophobicity, and clustering of 1.58. The low clustering gene Farsb has length 589amino acids, 31.6% hydrophobicity, and clustering of 0.69.

For randomly distributed amino acids of any length *N* and hydrophobicity *M*, this normalization makes the expectation of *ψ* equal to 1. For clustering at the nucleotide level, blocks of length *s* = 18 rather than 6 were used. Nucleotide clustering values were calculated for each possible permutation as to which nucleotides were scored as +1 and which as −1 (e.g. G and C as +1 and A and T as −1 constitutes one permutation). Amino acid clustering values *ψ* were Box-Cox transformed (λ=-0.29 for mouse, λ=-0.008 for yeast) prior to use in linear models, with mouse Q-Q plot shown in Figure S6B.

To generate a scrambled control sequence that is paired to each gene, we simply sampled its amino acids without replacement. To generate clustering-controlled scrambled sequences, 1000 scrambled sequences of each protein were produced, and the one that most closely matched the clustering value of the focal gene was retained. This left the average gene with a clustering value 0.0035 higher than its matched control, with the mean difference of the absolute deviation between a gene and its matched control equal to 0.0057, showing a close match with little directional bias. The mean value of each property was used across 50 scrambled sequences, but this led only to very modest reductions in confidence interval width relative to using a single scrambled control, e.g. ^~^20% in Figure 3. Because generating well-matched clustering-controlled scrambled sequences is computationally expensive, we used only a single matched-clustering scrambled control sequence per gene.

Transmembrane protein annotation was taken from both the “Membrane Proteins of Known 3D Structure” database (Stansfeld *et al.* 2015) (http://blanco.biomol.uci.edu/mpstruc/, including mouse proteins whose human homolog was experimentally verified as transmembrane, accessed July 16, 2017), and from UniProt (The UniProt Consortium 2017) (transmembrane annotation “experimental” ECO:0000269, “curated” ECO:0000303 and ECO:0000305, or “homology” to a “related experimentally characterized protein” ECO:0000250, accessed November 19).

## Data availability

Source data for the statistical analyses and figures are provided in Supplementary Tables S1-S6, available at Figshare and captioned in the main Supplementary Materials file. Code associated with generating and analyzing these tables is publicly available at https://github.com/MaselLab.

## Acknowledgements

This work was supported by the John Templeton Foundation (39667, 60814), and the National Institutes of Health (GM104040). The funders had no role in study design, data collection and analysis, decision to publish, or preparation of the manuscript. We thank Rafik Neme for insightful discussions, and Joost Schymkowitz and Rob van der Kant of the VIB Switch Laboratory for providing us with a stand-alone Waltz script.

## Author contributions

J.M. conceived the general approach, M.H.J.C. conceived the clustering metric, SLiM and transmembrane protein analyses, J.B. produced Figures 5 and S2, B.A.W. fitted statistical models and produced the other final figures, S.G.F. conducted all other upstream data analysis, and J.M. wrote the paper.

